# ECgo: All-Optical Induction of Single Endothelial Cell Injury and Capillary Occlusion in the Brain

**DOI:** 10.1101/2025.05.27.656398

**Authors:** Jacqueline Condrau, Srinivasa R. Allu, Tatiana V. Esipova, Eva Erlebach, Matthias T. Wyss, Chaim Glück, Luca Ravotto, Thomas Troxler, Marco Villa, Paola Ceroni, Mohamad El Amki, Sergei A. Vinogradov, Bruno Weber

**Affiliations:** Institute of Pharmacology and Toxicology, University of Zurich, Zurich 8057, Switzerland; Neuroscience Center Zurich, University and ETH Zurich, Zurich 8057, Switzerland; Department of Biochemistry and Biophysics, Perelman School of Medicine, University of Pennsylvania, Philadelphia, PA 19104, USA; Department of Chemistry, School of Arts and Sciences, University of Pennsylvania, Philadelphia, PA 19104, USA; G. Ciamician Department of Chemistry, University of Bologna, Bologna 40126, Italy; Department of Neurology, University Hospital Zurich, Zurich 8091, Switzerland

**Keywords:** Cerebral blood flow, brain capillaries, microstroke, porphyrins, photodynamic therapy

## Abstract

The ability to induce endothelial cell (EC) damage in the mouse brain with high spatial precision is invaluable for mechanistic studies of brain capillary injury and repair. Here, we introduce an optical method, termed ECgo, that utilizes a new two-photon-excitable porphyrin-based photosensitizer (*Ps2P*) to selectively obliterate single ECs within the brain microvascular network. Using the developed approach, we were able to induce occlusions of single capillaries with high spatiotemporal control, while preserving the surrounding tissue. Combined with longitudinal two-photon imaging, ECgo enables studies of morphological and functional consequences of targeted single capillary EC injury *in vivo* under healthy and diseased conditions.

**SIGNIFICANCE STATEMENT:** Brain capillary injury is a common feature of aging and many neurological disorders. While a single capillary lesion may appear inconsequential, the cumulative effect of repeated and spatially dispersed capillary insults can lead to substantial brain dysfunction. Understanding how single capillary injuries contribute cumulatively to long-term brain damage requires tools that can precisely target individual capillaries in the living brain. Here, we introduce an optical method that uses a new light-activatable compound to selectively injure single brain capillaries with high spatial accuracy. Our method enables detailed, longitudinal studies of capillary repair, blood flow recovery, local oxygen dynamics, and glial responses following microvascular injury.

## INTRODUCTION

Brain capillaries constitute over 80% of the cerebral vasculature and are essential for maintaining proper brain function. While occlusions of penetrating arteries and ascending veins can disrupt blood supply and homeostasis in large brain tissue regions^1–4^, the impact of single capillary injury and subsequent occlusion remains understudied. Notably, capillary occlusions have been observed in patients with cerebrovascular and neurological conditions, including vascular dementia^5^, cerebral microinfarcts^6, 7^, stroke^8, 9^, type 1 diabetes^10^, and Alzheimer’s disease^11, 12^, underscoring the need for better understanding their pathophysiology. Furthermore, microvascular pathology is hypothesized to both precede and contribute to age-related cognitive decline and neurodegenerative conditions^13, 14^.

Multiple experimental approaches have been used to induce capillary occlusions, including vascular injection of small occlusive microbeads^15^ and cholesterol crystals^16^, as well as optical methods, such as laser-induced injury and photothrombosis using Rose Bengal^1, 7, 17, 18^. However, these approaches lack the ability to create microvascular lesions with sufficient spatiotemporal accuracy, frequently leading to extensive non-local vascular injuries. Achieving precise, single-capillary occlusion in the living mouse brain remains technically challenging, hindering efforts to explore the molecular and cellular dynamics of capillary injury and repair across the interconnected neuron-glia-vasculature network.

Here, we report the development of a new technique, termed ECgo (Endothelial Cell guided obliteration), that allows targeting of single capillaries by inducing endothelial cell (EC) damage with high spatiotemporal control. ECgo makes use of a two-photon-excitable photosensitizer (termed *Ps2P*), based on the Zn complex of tetraarylphthalimidoporphyrin (TAPIP), which is closely related, both structurally and spectroscopically, to the two-photon oxygen probe Oxyphor 2P developed by us previously^19^. Unlike in disease-progressed states, where multiple capillaries are already damaged, ECgo makes it possible to create a single brain capillary occlusion to investigate its early impact, while avoiding the complexity of widespread vascular injury.

## RESULTS

### Development of the Two-Photon Photosensitizer *Ps2P*

Photosensitization processes and associated cell damage underpin photodynamic therapy (PDT) - an established clinical modality^20, 21^. The photodynamic effect (Fig. 1A) entails photoexcitation of a photosensitizer compound (Ps), usually to its lowest singlet excited state (S_1_), followed by a fast (pico-to-nanosecond) intersystem crossing (*isc*) with the formation of the Ps triplet state (T_1_). The T_1_ state is usually long-lived (microseconds), and, therefore, prior to its spontaneous deactivation (associated with rate constant *k* in Fig. 1A), it can undergo diffusional encounters and subsequent reactions with molecules in the environment. Direct interactions of the triplet with biological molecules (shown as RH in Fig. 1A; damaged species shown as RH*) fall under the umbrella of reactions classified as PDT Type I (electron transfer processes^22^). Since both the biomolecule and the sensitizer are typically large, slowly diffusing species, this mechanism usually requires close proximity of the Ps molecule to a biological target prior to photoexcitation.

**Figure 1:**
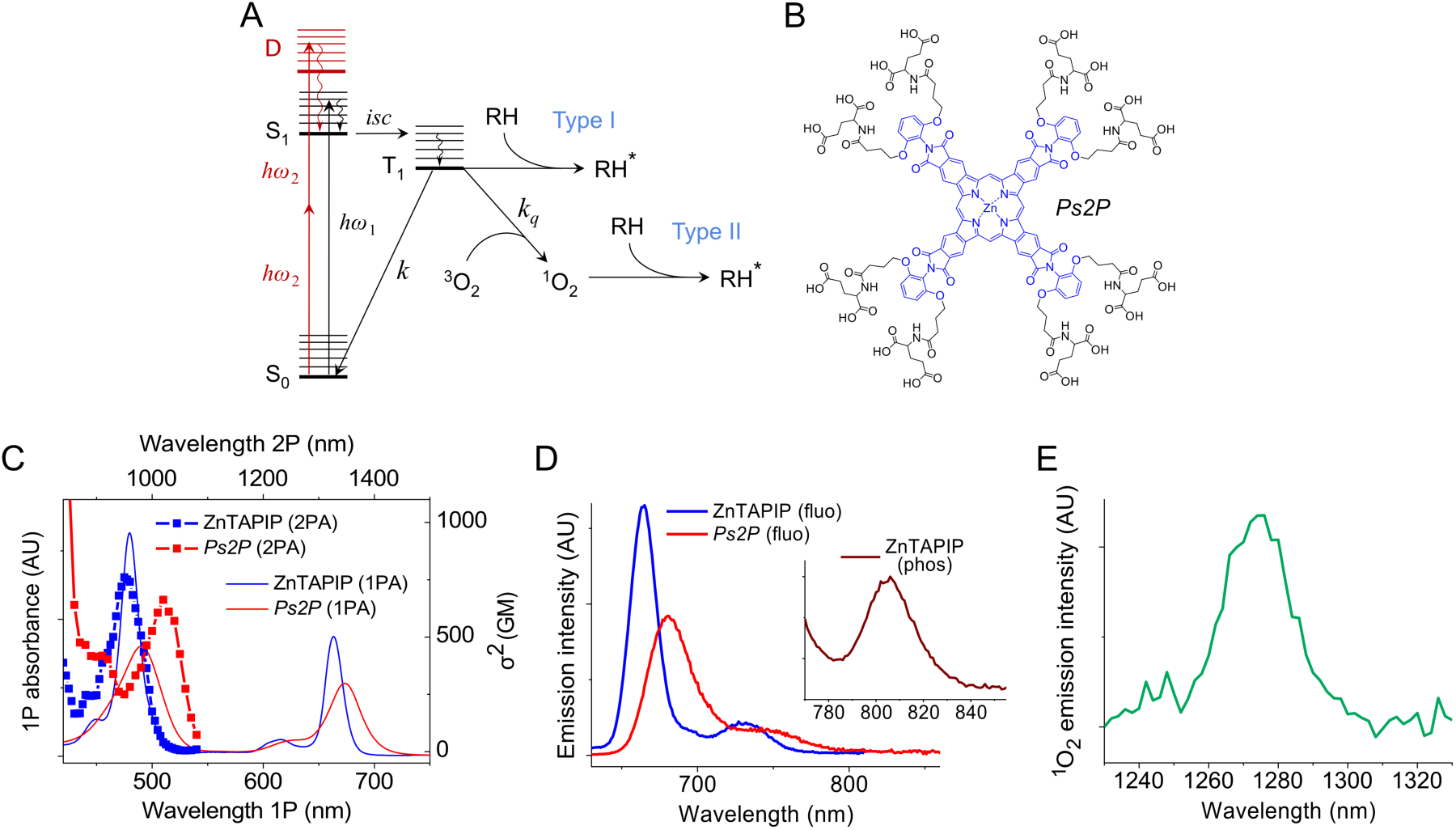
**(A)** Energy diagram showing one-photon (*hω*_1_) excitation of a photosensitizer to its lowest excited singlet state (S_1_), followed by intersystem crossing (*isc*) to the triplet state (T_1_), which reacts with biomolecules (RH) directly (Type I PDT), or sensitizes molecular oxygen, which subsequently reacts with biomolecules (Type II PDT), causing their damage. The 2P excitation pathway as shown (2ξ*hω*_2_; marked in red) is characteristic of porphyrins structurally similar to ZnTAPIP, in which the lowest two-photon active state Δ is located energetically above the Q state (S_1_), but below the B state. **(B)** Structure of the photosensitizer *Ps2P*. The parent Zn tetraarylphthalimidoporphyrin (ZnTAPIP) is shown in blue. **(C)** 1PA and 2PA spectra of ZnTAPIP in NMP (blue) and *Ps2P* (red). 1PA spectrum of *Ps2P* was recorded in aq. buffer (pH 7.3), while its 2PA spectrum was measured in D_2_O. **(D)** Fluorescence spectra of ZnTAPIP in aerated NMP (blue) and *Ps2P* in aerated aq. buffer, pH 7.3 (red) (λ_ex_ = 620 nm). The integrated emission intensities are normalized to unity. Inset: phosphorescence of ZnTAPIP in deoxygenated NMP. **(E)** Singlet oxygen emission sensitized upon irradiation of *Ps2P* in D_2_O at 475 nm.

Another more common mechanism, known as PDT Type II, involves energy transfer from the Ps triplet to the ground state molecular oxygen (^3^O_2_), resulting in the conversion of the latter to its singlet state. Singlet oxygen (^1^O_2_) is a rapidly diffusing, highly reactive species (lifetime ∼3.5 ms in H_2_O at ambient temperatures^23^) that can readily oxidize almost any molecule in the biological milieu. Thus, contributions of Type I via direct oxidation of organic targets and Type II pathways to the overall photodamage depend on the availability of O_2_, with Type II being the dominant pathway under normoxic conditions.

The combination of photosensitization with two-photon (2P) excitation gives rise to a 2P photodynamic effect^24, 25^, whereby 2P excitation makes it possible to confine photodamage to the immediate vicinity of the laser focus, permitting targeting of microscopic regions many microns deep in the tissue. For example, 2P excitation of a popular sensitizer called Rose Bengal has been used to create targeted damage in the brain^1, 3, 26^.

Porphyrins and phthalocyanines form a prominent group of photosensitizers due to their well-documented ability to generate triplet states^27^. However, thanks to their centrosymmetric electronic structures, regular cyclic tetrapyrroles typically possess extremely low two-photon absorption (2PA) cross-sections in the spectral region suitable for *in vivo* optical excitation. Much effort has been devoted over the years to the enhancement of 2PA in porphyrins and related compounds. A number of molecular systems have been developed that are able to act as efficient 2P absorbers and simultaneously singlet oxygen sensitizers. One relevant example is a porphyrin dyad with the peak 2PA cross-section (σ^2^) of ∼17,000 GM at 920 nm (Göppert-Mayer units: 1GM = 10^-50^ cm^4^s), which has been successfully used to occlude a single pial artery in the brain^28^.

In our previous work, we used aromatic π-extension combined with substitution by electron-withdrawing groups as a means to stabilize 2P-active *gerade* symmetry states (*g*-states) in porphyrins, creating compact, highly 2P-active porphyrinoids^29, 30^. The Pt(II) complex of one such porphyrin, Pt tetraarylphthalimidoporphyrin (PtTAPIP), was used to construct a high-performance two-photon oxygen probe Oxyphor 2P^19^. Due to the strong heavy metal-induced spin-orbit coupling, S_1_→T_1_ intersystem crossing (*isc*; Fig. 1A) in Pt and Pd porphyrins is extremely fast (picoseconds), and the triplet states are formed with nearly unity efficiency. However, the same spin-orbit coupling accelerates both radiative and non-radiative decays (rate constant *k* in Fig. 1A) of the triplet (T_1_→S_0_), effectively shortening its lifetime, which varies from tens to hundreds of microseconds (ms) for e.g. Pt and Pd TAPIPs, respectively^30^. We reasoned that using a lighter metal, such as Zn, could be beneficial for photochemical reactions involving the triplet state (both Type I and II pathways; Fig. 1A) since weaker spin-orbit coupling should elongate the T_1_ lifetime and thus increase the probability of diffusional encounters and subsequent photochemical transformations. Importantly, Zn is still able to promote efficient S_1_→T_1_ *isc* in porphyrins (albeit not as efficient as heavier metals) while not fully eliminating fluorescence, which is instrumental for imaging photosensitizer distributions in biological tissue.

The structure of the two-photon photosensitizer developed in this work, termed *Ps2P*, is shown in Fig. 1B. *Ps2P* is ZnTAPIP modified at the periphery with eight glutamic acid residues. The electronic absorption spectrum of the parent porphyrin, recorded in N-methylpyrrolidone (NMP), shows a familiar pattern consisting of two narrow transitions, Q (662 nm) and B or Soret (475 nm) bands, which are bathochromically shifted compared to the respective transitions of Pt and Pd TAPIPs^30^. The 16 carboxylate groups are able to solubilize *Ps2P* in aqueous media, although the corresponding absorption spectrum (Fig. 1C) clearly shows signs of aggregation, manifested by broadening and bathochromic shifts of the absorption bands (490 and 673 nm for B and Q bands, respectively) relative to the spectrum of the parent porphyrin (Fig. 1C).

The 2PA spectrum of *Ps2P* was measured in D_2_O, which has significantly weaker absorption near 900-1000 nm. The key spectral feature is the transition to a strongly 2P-allowed state (λ_max_∼1020 nm), which is reminiscent of the bands seen in the 2PA spectra of Oxyphor 2P^19^ as well as of parent Pt and Pd TAPIPs^30^. This *g*-symmetry state, termed Δ-state^31^, is forbidden for one-photon (1P) excitation, since the ground state in ZnTAPIP is also a *g*-state, and it is not seen in 1PA spectra. The energy of the Δ-state is below that of the 1P-allowed B (Soret) state. The strong 2PA to the blue of the Δ-peak allowed us to carry out efficient off-peak 2P photosensitization simultaneously with oxygen measurements using Oxyphor 2P without changing the laser wavelength. The 2PA maximum of Oxyphor 2P is near 960 nm, while at 1020 nm, its absorption is very low^19^.

Both *Ps2P* and its parent ZnTAPIP exhibit prompt and thermally activated (E-type) delayed fluorescence (TADF). In aerated NMP, ZnTAPIP fluoresces (λ_max_ = 665 nm) with a quantum yield (ϕ_fl_) of 0.14, showing a clear single-exponential fluorescence decay (𝜏_fl_ = 1.81 ns). Upon deoxygenation, the fluorescence quantum yield rises up to 0.46, and an additional band emerges (λ_max_ = 806 nm; Fig. 1D, inset), which corresponds to T_1_→S_0_ phosphorescence (ϕ_phos_∼0.015). The triplet lifetime obtained from either TADF (665 nm, S_1_→S_0_) or phosphorescence (806 nm, T_1_→S_0_) decays was found to be ∼9.4 ms, suggesting that the triplet of ZnTAPIP is efficiently quenched by O_2_. Indeed, no traces of the emission could be detected in air-saturated solutions.

The fluorescence quantum yield of *Ps2P* in aerated PBS was found to be reduced to 0.02, and the emission decay, which became profoundly non-single exponential, was considerably shortened (𝜏_av_ = 0.66 ns, λ_max_ = 695 nm). The reduction in fluorescence is likely caused in part by aggregation. Indeed, upon dissolving *Ps2P* in PBS containing bovine serum albumin (BSA; 4% by mass concentration), the spectral bands sharpened over time, and the fluorescence quantum yield increases by more than 3-fold, reaching ∼0.07, showing that BSA helps to prevent the aggregation of *Ps2P* and suggesting that in the blood *Ps2P* is predominantly bound to albumin and/or other hydrophobic/amphiphilic components. From a practical point of view, the fluorescence of *Ps2P* is highly instrumental for visualizing its distribution in the vasculature.

Upon deoxygenation, the fluorescence quantum yield of *Ps2P* (in PBS) increases by ∼10% (to ∼0.022), and a long-lived emission (detected near 700 nm) appears with a lifetime of ∼1.2 ms. While the phosphorescence of the *Ps2P* band could not be detected explicitly, the long-lived decay is most certainly a signature of its triplet state, which gives rise to the delayed fluorescence *via* reverse intersystem crossing (T_1_→S_1_).

To verify that *Ps2P* is capable of a Type II photodynamic action, ^1^O_2_ emission was recorded upon irradiation of a *Ps2P* solution at 475 nm (Fig. 1C). The quantum yield of ^1^O_2_ was found to be ∼0.2 (measured against standard sensitizer Eosin (ϕ(^1^O_2_) = 0.58)^32^. Given its relatively hydrophobic nature, *Ps2P* is likely to be bound to albumin and other proteins in the blood, but it could also be partially partitioned into the membranes of the ECs surrounding the capillaries. As such, upon photoexcitation, *Ps2P* could directly engage in photochemical processes with lipids and proteins in its vicinity according to the Type I mechanism.

### *Ps2P* Enables Targeted Capillary Occlusion in the Brain

Having characterized the chemical and photophysical properties of *Ps2P*, we next sought to evaluate its potential as a two-photon excitable photosensitizer for inducing single capillary occlusions in the mouse brain. To visualize the vascular network, we first intravenously injected the dye Texas Red-dextran (70 kDa, 2.5% in NaCl). Blood vessels were mapped through the cranial window at successive depths in the cortex, and the resulting images were used to construct a maximum intensity projection (MIP) image. Using the MIP image, several capillaries were selected for targeted occlusion based on their diameter (3-6 μm) and the depth from the surface (50-200 µm). In order to induce localized injury, *Ps2P* was injected intravenously (Fig. 2A) through a tail vein catheter (3 mg/ml PBS; estimated blood concentration 30 μM, assumed blood amount 0.078 ml/g body weight) and 2P-excited by scanning the laser focus along the vessel midline of a selected capillary (2-3 μm-long line, average power 30 mW under the objective, 960 nm), until blood flow ceased. Red blood cell (RBC) velocity (V_RBC_) in the capillary was measured before and after the irradiation scan (Fig. 2A-C). The first indication of the injury was detected as a slow-down and subsequent cessation of RBC passage through the capillary, occurring on average ∼60 s after the start of irradiation (Fig. 2A-C).

**Figure 2:**
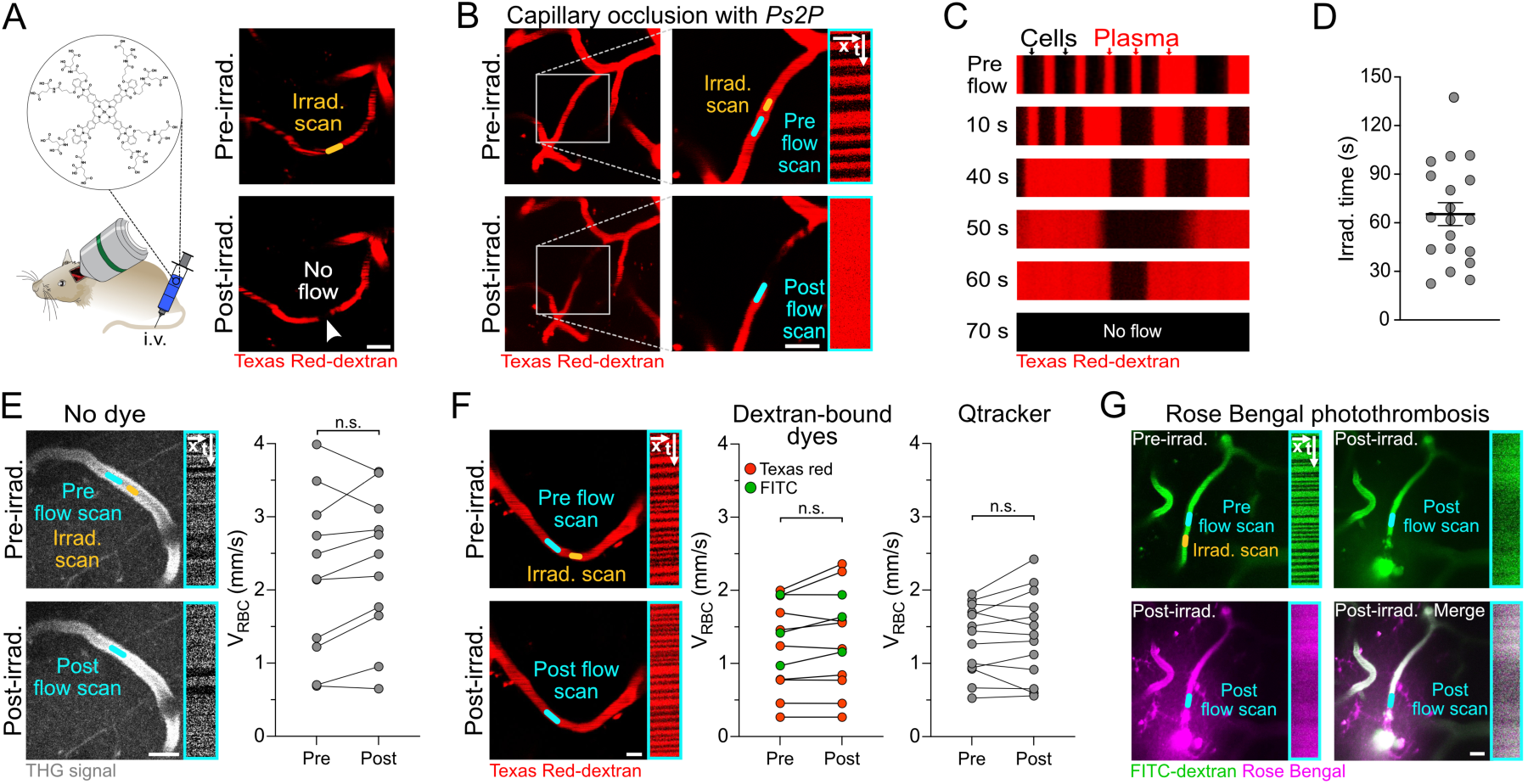
**(A)** Cartoon illustrating injection of *Ps2P* and representative two-photon fluorescence images of a capillary pre- and post-irradiation. The white arrowhead points at the occluded capillary segment. Scale bar: 10 μm. **(B)** Representative maximum intensity two-photon 30-µm-z-projections of a capillary pre-and post-induction of an occlusion with corresponding position of line scans (yellow = irradiation scan, cyan = flow scan). The line scan kymographs (shown within cyan rectangles, x = distance, t = time) confirm the presence (top) or absence (bottom) of blood flow. Scale bar: 10 μm. **(C)** Sequential line scan kymographs depicting a gradual reduction of blood flow to the point of full cessation at 70 s from the irradiation scan. **(D)** Duration of irradiation required to induce occlusion, with individual points representing individual capillaries (N = 3, n = 19). Mean ± SEM next **(E)** Left: Representative third harmonic generation (THG) images with the corresponding line scan kymographs (shown within cyan rectangles; x = distance, t = time) obtained in the absence of any dye. Scale bar: 10 μm. Right: No reduction in blood flow was detected (N = 3, n = 11). n.s. = not significant (two-tailed paired t-test). **(F)** Left: Control experiments using Texas Red-dextran (70 kDa) with the corresponding flow scans (shown within cyan rectangles, x = distance, t = time). Right: No reduction in blood flow was detected using different intravascular fluorescent dyes: red points = Texas Red-dextran (70 kDa; N = 2, n = 9), green points = FITC-dextran (70 kDa; N = 1, n = 3), Qtracker655™ (N = 4, n = 3). n.s. = not significant (two-tailed paired t-test). **(G)** Representative maximum intensity two-photon 20-µm-z-projections of a capillary irradiated in the presence of Rose Bengal, showing rupture and extensive dye extravasation. Corresponding line scan kymographs (right side, x = distance, t = time) indicate disrupted flow. Scale bar: 10 μm. N = number of animals, n = number of capillaries.

In the absence of *Ps2P* or other dyes, even prolonged irradiation (300 s; other irradiation parameters unchanged) did not lead to capillary occlusion and/or injury, as verified by label-free imaging using third-harmonic generation (THG) imaging (Fig. 2E)^33^. Similarly, irradiation in the presence of conventional intravascular dyes, e.g., Texas Red-dextran (2.5% in NaCl; 70 kDa), FITC dextran (2.5% in NaCl; 70 kDa), or quantum dot-based vascular dye (Qtracker655™; 2 μM), under identical conditions (300 s, 30 mW under the objective, 960 nm), resulted neither in a reduction in V_RBC_ nor in capillary stalling (25 capillaries tested; Fig. 2F). At the same time, induction of photothrombotic occlusion using Rose Bengal under comparable conditions (30 mW under the objective, 720 nm) led to a dramatic capillary insult, manifested by vessel rupture and leakage of the blood plasma and dye into the parenchyma (Fig. 2G). This is in stark contrast to occlusions using *Ps2P* (Fig. 2C-D), where the injury could be induced with high precision without vessel rupture.

### Irradiation in the Presence of *Ps2P* Leads to Blood Flow Stalling and EC Injury, but Not Clot Formation

Next, we sought to gain insight into the mechanism behind capillary blood flow stalls induced by photodynamic treatment using *Ps2P*. Vascular occlusions are commonly driven by (i) thrombus formation, particularly in large vessels, or (ii) stalling of blood cells in brain capillaries^8–10, 34^. First, we tested whether capillaries irradiated in the presence of *Ps2P* showed signs of platelet activation or fibrin aggregation. Animals were pretreated with anticoagulants, such as aspirin, heparin, or clopidogrel, before irradiation. Capillary occlusion occurred in 90% of aspirin-, 94% of heparin-, and 100% of clopidogrel-treated capillaries (Fig. 3A-B). Furthermore, we tested whether a recombinant tissue plasminogen activator (rtPA), an FDA-approved reperfusion drug, injected 30 min after the occlusion, could restore blood flow. In all cases, rtPA did not lead to capillary recanalization, confirming that capillary stalls in our model are not due to fibrin-driven intravascular clot formation (Fig. 3A-B).

**Figure 3:**
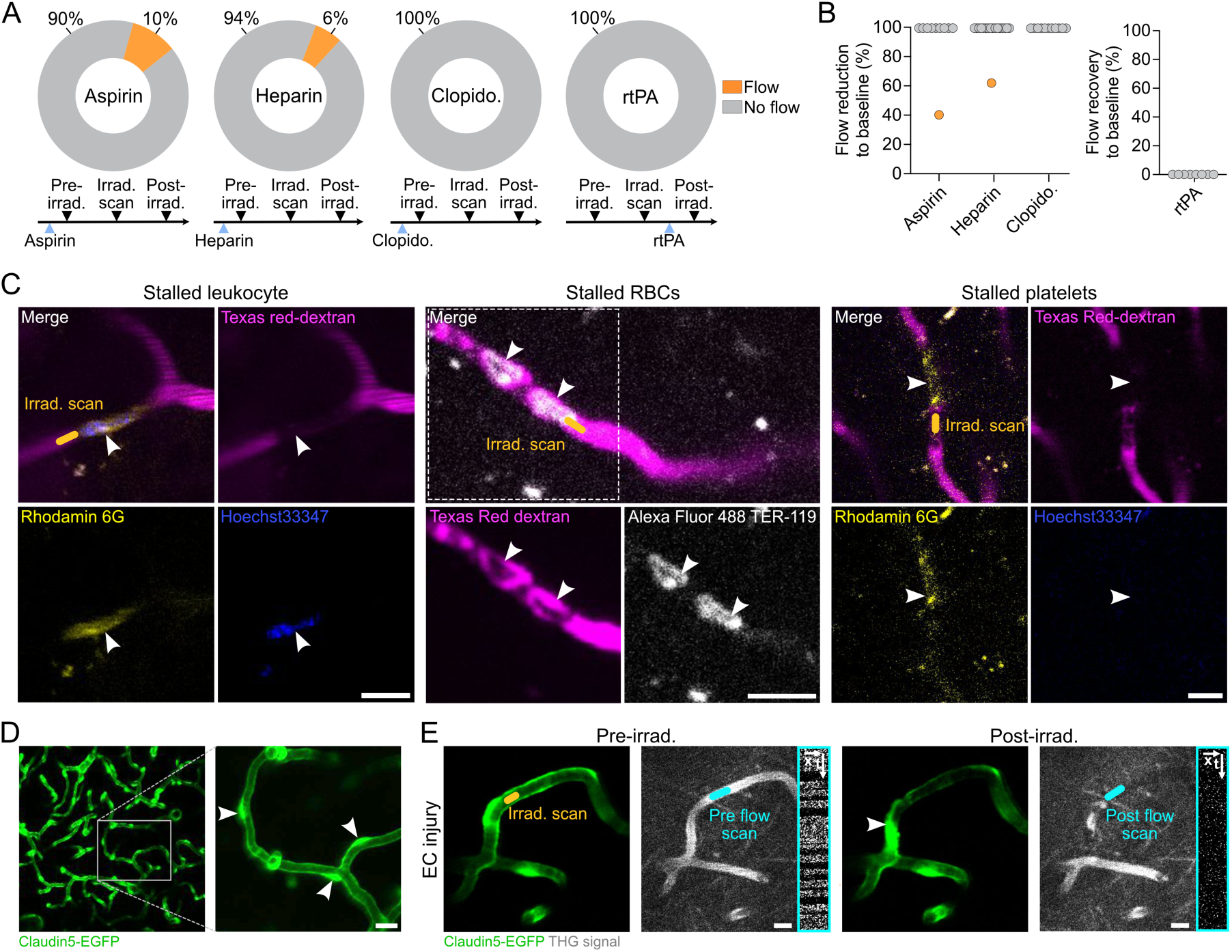
**(A)** Quantification of patent capillaries subjected to irradiation in the presence of *Ps2P*, either pretreated with anticoagulants to prevent occlusion or post-treated with rtPA to promote recanalization. All treatments failed to prevent blood flow cessation or to restore blood flow, respectively. **(B)** Reduction in blood flow upon capillary irradiation in the presence of anticoagulant drugs (left): Aspirin (N = 2, n = 10), heparin (N = 3, n = 17), clopidogrel (N = 4, n = 12) or blood flow recovery following reperfusion therapy with rtPA (N = 2, n = 18; right). Points represent single capillaries. **(C)** Representative two-photon images showing stalled blood cells inside the targeted capillary (left: leukocyte, middle: red blood cells (RBCs), right: platelets). Channels are displayed separately, with the merged image in the top left. Scale bars: 10 μm. **(D)** Representative maximum intensity two-photon 30-µm-z-projections of a Claudin5-EGFP reporter mouse. White arrowheads indicate individual EC nuclei. Scale bar: 10 μm. **(E)** Representative two-photon images of an EC injured capillary, showing pre-irradiation EC morphology (left) and post-irradiation damage (right). Line scan kymographs (cyan rectangles) using third-harmonic generation (THG) imaging confirm the presence and absence of blood flow pre- and post-irradiation, respectively. White arrowhead points towards injured EC nuclei. Scale bars: 10 μm. N = number of animals, n = number of capillaries.

Next, we tested whether capillary stalls were caused by a specific type of transiting cells. Leukocytes, platelets, and red blood cells (RBCs) were labeled using well-established cellular markers Rhodamine 6G, Hoechst 33342, and Alexa Fluor 488 TER-119 (Fig. 3C). Leukocytes exhibited dual staining with Rhodamine 6G and Hoechst 33342^8^; platelets were stained with

Rhodamine 6G only, and RBCs were labeled with Alexa Fluor 488 TER-119. In addition, Texas Red-dextran was injected to label the blood plasma. Subsequent capillary irradiation in the presence of *Ps2P* led to clogging of capillaries by individual cells, including leukocytes, RBCs, or platelets. However, the stalling cells were random, and no specific cell type could be identified as the cause of the occlusion. Instead, upon irradiation and the resulting decrease in blood flow, transiting cells became lodged in the capillary segment adjacent to the irradiation site, contributing to the resulting capillary occlusion.

Having demonstrated that capillary occlusion in our model does not result from classical intravascular coagulation pathways, we turned our attention to vessel wall injury, specifically endothelial injury, as a potential underlying mechanism. Using Claudin5-EGFP endothelial reporter mice, we analyzed EC morphology before and after irradiation. Indeed, we observed EC damage at the site of irradiation in the targeted capillary (Fig. 3D-E), suggesting that the capillary occlusion is likely caused by local EC injury rather than by intravascular clot formation or blood cell stalling.

### Consequences of Single Capillary Injury: Capillary Recanalization, Changes in Local Hemodynamics, pO_2_, and Microglial Activation

To study the fate of the damaged capillaries, we performed longitudinal two-photon imaging of the adjacent microvascular network directly after induction of occlusion, as well as 24 h and 48 h post-irradiation. Remarkably, blood flow was restored in all targeted capillaries 24-48 h post-irradiation, revealing a rapid brain capillary repair response (Fig. 4A-B).

**Figure 4:**
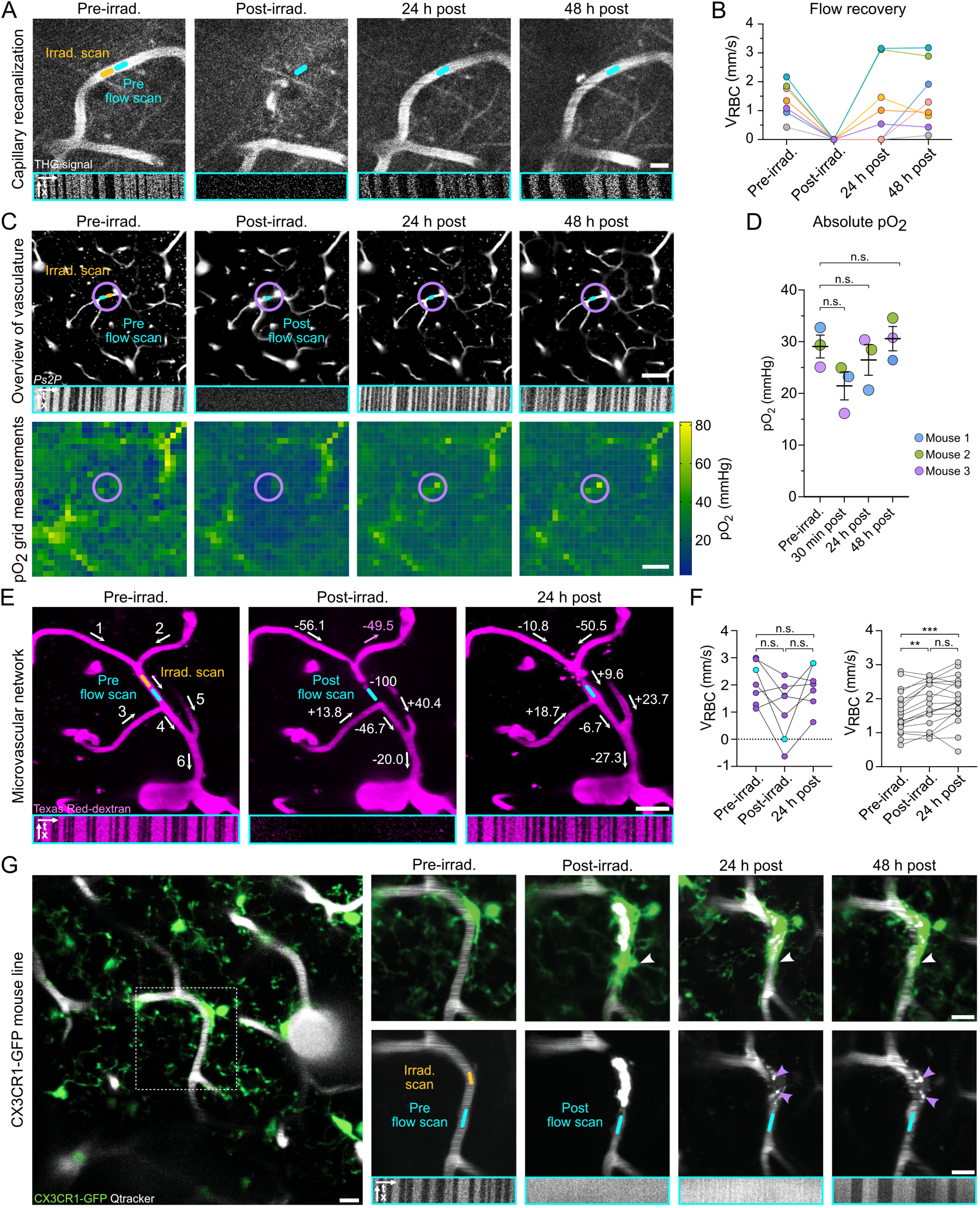
**(A)** Representative two-photon images showing capillary recanalization 24 h post-irradiation with corresponding line-scan kymographs (cyan rectangle, bottom). THG = third-harmonic generation. **(B)** Quantification of V_RBC_ recovery 24 h and 48 h post-irradiation. Each point represents an individual capillary (N = 3, n = 8). **(C)** Local tissue partial pressure of oxygen (pO_2_) pre- and post-irradiation. Top panels: Two-photon images of the region of interest surrounding the occluded capillary recorded pre-irradiation, immediately post-irradiation, as well as 24 h and 48 h post-irradiation, with the corresponding line-scan kymographs (shown within cyan rectangle). Bottom panels: 28 × 28 grid-like measurements of tissue pO_2_ in awake mice performed pre-irradiation, immediately post-irradiation, as well as 24 h and 48 h post-irradiation. The magenta circle (50 µm radius) marks the region quantified in *(D)*. Scale bars: 50 μm. **(D)** Absolute pO_2_ values within the circular region shown in *(C)*. Each data point represents a single capillary occlusion (one occlusion per mouse), where different colors correspond to individual animals (N = 3, n = 3). Mean ± SD, p > 0.05, Friedman test. **(E)** Local blood flow changes induced by single capillary occlusion. Maximum-intensity two-photon 30 µm z-projections with the corresponding line-scan kymographs (shown within cyan rectangles, bottom). Numbers indicate neighboring capillary segments. Relative RBC velocity (V_RBC_) changes in neighboring capillaries (1-5) post-irradiation. Scale bar: 10 μm. **(F)** Left: Quantification of V_RBC_ in capillaries pre- and post-irradiation from the example in (E) (N = 1, n = 7). Right: Quantification of V_RBC_ in the adjacent microvacuolar network (N = 1, n = 26). The occluded capillary is shown in cyan. Magenta points represent adjacent capillaries (capillaries 1-6 in (E)), while gray points correspond to capillary segments in the surrounding microvascular network that are not directly attached to the occluded capillary and not shown in the z-stack projection. *p < 0.05, repeatedly measured one-way ANOVA. Mean ± SEM. ***(G)*** Microglial response to capillary occlusion. Top panels: local microglial activation post-irradiation in CX3CR1-GFP reporter mouse, with microglial processes ensheathing the injured capillary (white arrowheads). Bottom panels: images of the fluorescence of an intravascular dye (Qtracker655™ pre- and post-irradiation, Texas Red-dextran 24 h and 48 h post-irradiation), showing the presence or absence of the blood flow in the targeted capillary pre- and post-irradiation. 48 h after irradiation, the capillary is reinstated. Magenta arrowheads indicate internalized vascular dye. Scale bars: 10 μm. N = number of animals, n = number of capillaries.

Capillary flow stalls may lead to changes in perfusion, oxygenation, and neuroinflammation in the affected area. To date, research on brain capillary stalls and injury has primarily focused on multiple and widespread capillary occlusions, while the effects of single capillary occlusion remain largely unknown. ECgo provides a valuable tool to manipulate one brain capillary at a time in order to assess local hemodynamics, tissue oxygenation, and microglial responses.

To evaluate the impact of capillary occlusion and subsequent repair on local tissue oxygenation, we used Oxyphor 2P, a previously developed pO₂ sensor^19^, and two-photon phosphorescence lifetime microscopy^35, 36^ to image the pO_2_ distribution in the area adjacent to the occlusion in awake animals, as described previously^37^. pO_2_ in the surrounding tissue was measured prior to the induction of occlusion (baseline pO_2_), 30 min post-irradiation, 24 h, and 48 h post-irradiation (Fig. 4C). Within the examined two-dimensional circular region (50 µm in diameter) surrounding the occlusion (Fig. 4D), mean tissue pO_2_ decreased by 26.6% (from 29.1 mmHg to 21.4 mmHg) immediately after the occlusion but returned to the baseline level within 24-48 h upon blood flow recovery (Fig. 4D). Notably, the observed pO_2_ drop did not reach hypoxic levels (<10 mmHg), indicating that the surrounding microvascular network was able to compensate for the local damage and maintain oxygen delivery to tissue.

Next, we examined how the occlusion of a single capillary impacts hemodynamics in the surrounding capillary network. A selected capillary was irradiated, and V_RBC_ was measured in the adjacent capillaries. Following the occlusion, the direction of blood flow changed such that the flow in one of the feeding capillaries, located upstream from the occluded capillary, increased (capillary 2 in Fig. 4E). The analysis of the flow in the wider capillary network surrounding the occlusion revealed that overall V_RBC_ in the affected area increased (Fig. 4F), suggesting that upon a single capillary occlusion, the collateral flow compensates for the local perfusion deficit. Importantly, our data show that while blood flow ceased in the targeted capillary, neighboring vessels remained patent, highlighting the robustness of the cortical capillary network and suggesting the presence of a mechanism that signals the adjacent network to compensate for single capillary occlusion.

Local microglial cells are known to react rapidly to local cell damage^38–40^. To test whether capillary injury in our model led to a local inflammatory response and subsequent microglial activation, we used CX3CR1-GFP transgenic reporter mice. We observed that microglia became locally activated after capillary injury, extended their processes to ensheathe the injured capillary, and internalized the leaked vascular probe Qtracker655™, as indicated by co-expression of vascular and GFP signals (Fig. 4G). This microglial activation was seen both in cells directly next to the occluded capillary (Fig. 4G) and in nearby microglia that were not in direct contact with it, suggesting that local inflammatory signaling occurs in response to EC injury.

## DISCUSSION

Here, we have introduced a new porphyrin-based photosensitizer, *Ps2P*, and demonstrated its use for induction of single-EC injury and capillary occlusion in the living brain under 2P excitation, an approach we termed ECgo. Compared to methods employing conventional photosensitizers, ECgo is characterized by greater precision, lower off-target damage, and overall, appears to have broader applicability for longitudinal studies of microvascular dysfunction. The most important feature of ECgo is that it allows for a highly accurate single capillary occlusion, avoiding vessel rupture and collateral damage to neighboring vessels and tissue. Interestingly, we report that the damaged capillaries undergo fast repair and regain blood flow within 24-48 h after injury. Detailed investigation of this remarkable endothelial repair is the focus of a separate study^41^. Thus, ECgo provides a new, useful tool for studying the effects of single capillary injury on local blood flow, oxygenation, and neuroinflammatory responses.

Photodynamic methods have been used previously to create capillary occlusions. Perhaps the most known photosensitizer is Rose Bengal^42–44^, which is widely used for producing large vessel occlusions^1, 3^. However, the use of Rose Bengal is associated with thrombosis and is often followed by vessel rupture even upon focused 2P-excitation of the sensitizer^43^. In contrast, occlusion with *Ps2P* did not lead to thrombosis or cause rupture. In fact, our results indicate that the injury could be localized to the endothelium, such that the passage of blood cells was impeded. In our hands, using the photosensitizer Rose Bengal, it was not possible to create such precise single-capillary injuries. In this regard, a pertinent question arises: what properties of *Ps2P* are responsible for its unique ability to induce such a precise single EC injury? A separate study would be required to delineate the exact mechanism of action of *Ps2P*; however, its relatively large aromatic structure and resulting hydrophobicity suggest that it could be partially distributed into the membranes and/or interiors of ECs that make up capillary walls. This contrasts with smaller, more hydrophilic sensitizers (e.g., Rose Bengal), which are presumably localized to the vascular lumen. Once associated with an EC and subjected to 2P excitation, *Ps2P* can operate *via* Type I and/or Type II photodynamic mechanisms (Fig. 1), leading to selective destruction of the targeted cell. Importantly, the excitation volume in which the triplet state of *Ps2P* is produced and engages in subsequent photochemistry can be adjusted by regulating the photon flux^45^, such that at mild excitation fluxes, the damage can be confined to a single EC. On the other hand, ^1^O_2_ generated by *Ps2P* within the capillary lumen is expected to be scavenged by albumin molecules, to which *Ps2P* is likely to be bound. These partially damaged albumin molecules would be carried away and dispersed by the blood flow, not leading to the formation of a clot. The above scenario is speculative at this point, but it suggests one potentially testable hypothesis behind the highly selective action of *Ps2P*.

Capillary stalls are common in many cerebrovascular and neurodegenerative diseases^6, 8, 9, 11, 12^. However, the development of reliable and precise techniques for inducing targeted single-capillary stalls in the rodent brain has remained an unmet challenge. Current approaches to induce capillary occlusion rely on the use of small occlusive materials, such as various polymeric microbeads or cholesterol crystals^15, 16^. However, such methods do not permit targeting of a single capillary in a pre-selected location. In contrast, the two-photon photodynamic approach using *Ps2P* makes it possible to induce occlusions one at a time at desired cortical depths while avoiding widespread clot formation and/or systemic vascular damage. These properties constitute a distinct advantage of ECgo compared to conventional methods.

From a hemodynamic point of view, single capillary occlusions have been previously modeled *in silico*^46^, demonstrating that changes in local capillary flow depend on baseline flow and local capillary network topology. Occlusion of a capillary segment with two inlet and two outlet vessels has the most severe impact on local tissue perfusion. In that case, blood flow is reduced by more than 30% in capillaries located within two branching orders from the occlusion site^46^. Our current experimental results are in line with these modeling studies, showing that a single capillary occlusion causes redistribution of flow, suggesting that oxygen delivery is compensated by the local microvascular network. Indeed, while occlusion of a single capillary led to a drop in tissue oxygenation in the area surrounding the capillary, pO_2_ did not reach hypoxic levels. However, occlusion of multiple capillaries could cause local hypoxic regions, resulting in tissue damage. However, how these microvascular events might contribute to long-term cerebrovascular impairment merits further investigation.

Single capillary occlusions contribute to neurodegenerative and cerebrovascular diseases, where progressive capillary dysfunction leads to chronic perfusion deficits^14, 47–49^. Capillary occlusions and flow stops have been observed in aging^13^, Alzheimer’s disease^12, 49^, Type 1 diabetes^10^, and vascular dementia^5^. Employing ECgo, it is possible to create isolated capillary occlusions in an otherwise intact vascular network, offering a precise tool for investigating early microvascular dysfunction before the onset of widespread damage. Such studies are important to advance our understanding of the vascular contribution to Alzheimer’s disease and vascular dementia, where microvascular damage is thought to precede overt neurodegeneration^47, 49, 50^. Little is known about cellular and molecular responses to EC injury in the intact brain. Capillary repair and EC responses may vary across brain regions and conditions. Future studies will examine the impact of single capillary occlusion in different brain regions, aged animals, and in disease states affecting the cerebrovasculature. Our preliminary results suggest that damaged capillaries exhibit rapid repair, though the mechanisms driving this process are still unclear. ECgo offers a valuable tool for studying EC repair across pathophysiological contexts where maintaining vascular integrity is critical.

## MATERIALS AND METHODS

### Ps2P Synthesis and Photophysical Experiments

#### General Information

All solvents and reagents were purchased from standard commercial sources and used as received. NH_2_GluOAll‧TsOH^51^ and H_2_TAPIP-OH^30^ were synthesized as described previously. Thin-layer chromatography was performed on aluminum-backed silica gel TLC plates (SiliaPlate™ 200 μm, with F254 indicator, Silicycle). Column chromatography was performed on silica gel (Silicycle Silica Flash F60, 60 Å, 40-63 µm 230-400 Mesh) using air-forced flow (0.5-1.0 bar). Flash chromatography was performed using an automated system (Combiflash *NextGen* 300, Combiflash) equipped with a UV-Vis detector. ^1^H, ^13^C, ^19^F NMR spectra were recorded on Bruker NEO 400 (operating at 400 MHz for ^1^H, 101 MHz for ^13^C), Bruker DRX 500 (operating at 500 MHz for ^1^H, 126 MHz for ^13^C), or NEO 600 (operating at 600 MHz for ^1^H, 150 MHz for ^13^C) spectrometers. Deuterated solvents (CDCl_3_, DMSO-*d*_6_) were purchased from Cambridge Isotope Laboratories, Inc. with TMS as an internal standard. NMR data were analyzed using MestReNova software. Mass spectra were recorded on a MALDI-TOF Bruker Daltonics Microflex LRF instrument, using 1,8,9- trihydroxyanthracene (dithranol) or α-cyano-4-hydroxycinnamicacid (CCA) as a matrix in positive-ion or negative-ion mode.

Optical spectra were recorded on a Perkin-Elmer 365 UV-Vis spectrophotometer. Steady-state fluorescence measurements were performed on an FS920 spectrofluorometer (Edinburgh Instruments, UK), equipped with R2658P red-sensitive PMT (Hamamatsu). Quartz fluorometric cells (1 cm path length; Starna) were used in all optical experiments. For quantum yield measurements, the absorbances of the samples at the excitation wavelengths were kept below 0.05 OD. The excitation and emission slits in the fluorometer were 2 nm. The quantum yield measurements were performed according to the published guidelines relative to the fluorescence of Oxazine 1 (ϕ_fl_ = 0.15, 22 °C, EtOH)^52^.

Time-resolved fluorescence measurements were performed using a time-correlated single photon counting (TCSPC) system consisting of a picosecond diode laser (PicoQuant), MCP-PMT detector (Hamamatsu R2809U) and a TCSPC board (Becker & Hickl, SPC-730).

The two-photon absorption (2PA) spectra were measured by the two-photon fluorescence excitation method against fluorescence of Rhodamine B (σ^2^ (800 nm) = 180 GM)^53^, which was used as a standard. The setup for 2PA measurements was described previously^29^.

#### H_2_TAPIP(GluOAll)_8_

To a solution of H_2_TAPIP-OH (0.020 g, 0.011 mmol) in DMF (3 ml), HBTU (0.064 g, 0.168 mmol) was added, and the reaction mixture was stirred at room temperature (r.t.) for 10 min (Supplementary Fig. S1). N,N-diisopropylethylamine (0.088 ml, 0.504 mmol) and NH_2_GluOAll·TsOH (0.067 g, 0.168 mmol) were added to the mixture, and the resulting solution was stirred at r.t. for 5 days. The reaction mixture was poured into an ice-cold aqueous solution of HCl (10%, 20 ml), and the green precipitate was collected by centrifugation, washed with water (2 × 10 ml) and dried in vacuum. The product was purified by silica gel column chromatography using dichloromethane (DCM):methanol (100:3) mixture as an eluent. The target compound was eluted as a green band, which was collected. The solvent was removed by rotary evaporation, and the product (green solid) was dried in vacuum. Yield: 0.03 g (81%). ^1^H NMR (DMSO-*d_6_*, 80 °C), δ (ppm): -2.10 (s, 2H), 1.87-1.94 (m, 32H), 2.14-2.25 (m, 32H), 4.15-4.22 (m, 24H), 4.42-4.54 (m, 32H), 4.96-5.09 (s, 32H), 5.63-5.74 (m, 14H), 6.94 (d, J = 8.65 Hz, 8H), 7.52 (t, J = 8.55 Hz, 4H), 7.83 (d, J = 6.45 Hz, 8H), 10.77 (s, 8H), 12.07 (s, 4H) (Supplementary Fig. S2).

#### ZnTAPIP(GluOAll)_8_

To a solution of H_2_TAPIP(GluOAll)_8_ (0.03 g, 0.0084 mmol) in DMF (8 ml), an excess of Zn(OAc)_2_ ‧2H_2_O (20 eq, 0.037 g, 0.168 mmol) was added, and the mixture was refluxed at 130 °C for 1 h. The reaction progress was monitored by measuring UV-vis absorption spectra. The mixture was cooled to r.t. and poured into ice-cold water. The resulting green precipitate was collected by centrifugation and dried in vacuum. Crude ZnTAPIP(GluOAll)_8_ was introduced into the next step without further purification.

#### Ps2P (ZnTAPIP(GluOH)_8_)

ZnTAPIP(GluOAll)_8_ (0.018 g, 0.0049 mmol) was dissolved in DMF (2 ml). Pd(PPh_3_)_4_ (0.046 g, 0.0395 mmol) and morpholine (0.34 ml, 3.92 mmol) were added, and the mixture was stirred for 48 h at r.t. The solvents were removed in vacuum, THF (10 ml) and acetic acid (0.5 ml) were added, and the resulting suspension was sonicated on an ultrasound bath (Branson) for 10 min. The title compound was isolated by centrifugation as a green solid, washed with THF and then twice with Et_2_O, and dried in vacuum. Yield: 0.014 g, 93%. ^1^H NMR (DMSO-*d_6_*, 80 °C), δ (ppm): 1.89-1.97 (m, 32H), 2.18-2.27 (m, 32H), 3.58-3.61 (m, 32H), 4.15-4.20 (m, 24H), 6.93 (d, J = 8.40 Hz, 8H), 7.50 (t, J = 8.15 Hz, 4H), 7.72-7.56 (m, 16H), 7.83-7.89 (m, 16H), 10.69 (s, 8H), 11.88 (s, 4H), 14.66 (s, 16H) (Supplementary Fig. S3).

#### Singlet Oxygen Sensitization

The quantum yield of singlet oxygen (^1^O_2_) generation by *Ps2P* was determined relative to that by Eosin Y: ϕ(^1^O_2_)_Eosin_=0.58^32^. Air-equilibrated isoabsorbing solutions of *Ps2P* and of the standard (Eosin) were excited at the 475 nm, and ^1^O_2_ phosphorescence was detected in the 1230–1330 nm range. For each measurement, four spectra were averaged. The singlet oxygen generation quantum yield was calculated using the following formula:

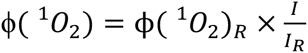

I is the integrated intensity of the phosphorescence of ^1^O_2_ sensitized by *Ps2P*, and *I_R_* refers to the ^1^O_2_ phosphorescence sensitized by the standard.

### Ethical Approval and Animal Welfare

This study was conducted following the Swiss Animal Protection Law and approved by the Swiss Veterinary Office, Canton of Zurich (Animal Welfare Act of December 16, 2005, and Animal Protection Ordinance of April 23, 2008). The local Cantonal Veterinary Office in Zurich granted ethical approval under licenses ZH169/2017 and ZH152/2021.

### Laboratory Animals

This study utilized male and female mice between the ages of two and eight months. The animals were housed under a reversed 12:12 h light-dark cycle with unrestricted access to food and water. Standard laboratory chow was provided unless stated otherwise. Wild-type and control animals were C57BL/6J mice obtained from Charles River. For the microglia studies, we employed CX3CR1-GFP mice (B6.129P-Cx3cr1/J; Jackson, Stock No.: 005582; Gensat.org). Investigations on ECs utilized Claudin5-EGFP mice (B6.Cg-Tg(Cldn5-EGFP)Cbet/U).

### Head Plate Surgery

Thirty minutes before the surgical procedure, mice received subcutaneous buprenorphine (0.1 mg/kg; Temgesic; Indivior Schweiz AG). Anesthesia was induced with 5% isoflurane and maintained at 1.5% via a face mask, with a flow rate of 300 ml/min of oxygen and 600 ml/min of compressed air. A heating mat was used to maintain the body temperature at 37 °C. VitA Pos (Bausch + Lomb) eye ointment was applied to prevent corneal dehydration. The scalp was shaved, cleaned, and disinfected with Kodan (Schülke & Mayr AG) before securing the animal’s head in a stereotactic frame (Model 900; David Kopf Instruments). A midline scalp incision (∼1.5 cm) was made, and the periosteum was removed. A bonding agent (Gluma Comfort Bond; Heraeus Kulzer) was applied and polymerized using blue light (600 mW/cm²; Demetron). Using a light-curing composite, a stainless-steel head plate was affixed centrally on the skull, avoiding the left somatosensory cortex region for craniotomy (Synergy D6 Flow; Coltene AG). The wounds were sealed with acrylic glue (Histoacryl; B. Braun) and antibiotic ointment (Neomycin; Cicatrex; Janssen-Cilag AG). To prevent dehydration, 10 ml/kg of pre-warmed ringerfundin was administered intravenously.

### Craniotomy and Cortical Window Implantation

Mice were anesthetized using a subcutaneous injection of fentanyl (0.05 mg/kg; Sintenyl; Sintetica), medetomidine (0.5 mg/kg; Domitor; Orion Pharma), and midazolam (5 mg/kg; Dormicum; Roche). The body temperature was maintained at 37 °C using a heating mat, and 100% oxygen was supplied at 300 ml/min via a face mask to prevent hypoxia. A 4x4 mm craniotomy was performed over the left somatosensory cortex using a dental drill (H-4-002HP; Rotatec GmbH). A 3 × 3 mm sapphire window (Powatech GmbH) was then placed over the brain surface and immobilized with a custom fixture, secured with a bonding agent (Gluma Comfort Bond; Heraeus Kulzer) and light-curing composite (EvoFlow; Ivoclar). Postoperative care encompassed subcutaneous injections of buprenorphine (0.1 mg/kg), carprofen (10 mg/kg; Rimadyl; Pfizer), ringerfundin (10 ml/kg), and an antagonist mix of flumazenil (0.5 mg/kg; Anexate; Roche) and atipamezole (2.5 mg/kg; Antisedan; Pfizer). Mice were permitted to recover for a minimum of two weeks prior to imaging.

### *In Vivo* Two-Photon Imaging

All two-photon imaging experiments were performed using a custom-designed two-photon laser scanning microscope^54^. The system incorporated a femtosecond-pulsed laser (Chameleon Discovery TPC; Coherent/Spectra Physics) with a tunable excitation wavelength range of 680– 1300 nm, a pulse duration of approximately 120 fs, and an 80 MHz repetition rate. Imaging was conducted using either a 25× water-immersion objective (W Plan-Apochromat 25×/1.05 NA; Olympus) or a 20× water-immersion objective (W Plan-Apochromat 20×/1.0 NA; DIC VIS-IR; Zeiss). Emitted fluorescence was separated from the excitation path using an 825 nm short-pass (SP) dichroic mirror and further divided into four detection channels using additional dichroic mirrors (506, 560, and 652 nm). Each channel was equipped with specific short-pass and band-pass filter combinations: CH1 (770 SP + 475/64 BP), CH2 (770 SP + 535/50 BP), CH3 (770 SP + 607/70 BP), and CH4 (810 SP + 824 SP + 990 SP). Fluorescence signals were detected and amplified using photomultiplier tubes (PMTs; CH1–3: H10770PA-40 SEL, Hamamatsu; CH4: H10770PA-50 SEL, Hamamatsu). PMT gain was regulated via custom software written in LabVIEW. Image acquisition and system control were managed using a customized version of ScanImage r3.8.1^55^.

### Imaging of Cerebral Vasculature

Animals underwent head plate surgery and chronic cranial window implantation to enable head fixation for subsequent two-photon imaging (see Supporting Information for details). The visualization of cortical blood vessels was achieved through the intravenous administration of fluorescent tracers. The choice of tracer depended on the mouse strain and its ability to express fluorescent proteins. Specifically, 50 μl of Texas Red-dextran (2.5% in NaCl; 70 kDa; Thermo Fisher Scientific, D1830), FITC Dextran (2.5% in NaCl; 70 kDa; Sigma; 46945-100MG-F), or 20 μl of Qtracker655™ (Thermo Fisher Scientific, Q21021MP) was injected.

### Blood Flow Measurements

The assessment of blood flow velocity (V_RBC_) was performed using line scans. Red blood cell velocity was determined by performing a line scan along the vessel’s midline (256 × 256 pixels; 0.64 ms per line; recorded over 72 frames, corresponding to 11.8 s). Subsequent analysis of the line scans was performed using a custom image processing toolbox (Cellular and Hemodynamics Processing Suite^56^) in conjunction with the RADON^57^ transform in MATLAB (R2017b; MathWorks).

### Drug Application

Intravenous injection of 0.2 IU/g Heparin (Heparin-Na; B. Braun) was performed via the tail vein a few minutes before capillary irradiation. Clopidogrel (Tocris, Cat.# 1820) administration included either a 4-day treatment at a lower dose (20 mg/kg; one i.p. injection per day over 4 d) or a higher single dose injection (100 mg/kg, i.p.) on the capillary irradiation day, with a minimum 3-h waiting period between injection and capillary irradiation. Aspirin application (100 mg/kg; Sigma, A5376) was performed through either i.p. or i.v. injection 3 h prior to capillary irradiation. For rtPA (Human t-PA; Actilyse; Boehringer Ingelheim) administration, intravenous injection via the tail vein was performed post-irradiation (10 mg/kg; 10% bolus; 90% perfusion; i.v.)^58^. After starting the rtPA infusion, the targeted vasculature was monitored for at least 125 min to assess recanalization.

### *In Vivo* Staining of Blood Cells

Cells within blood vessels were identified based on their morphology and staining patterns using Rhodamine 6G and Hoechst 33342. Blockages caused by red blood cells (RBCs) appeared as dark, hollow areas, while platelets were observed as clusters of small green particles. Neutrophils were the only cells that exhibited double staining with Rhodamine 6G and Hoechst 33342^8^.

### Photothrombosis using Rose Bengal

Rose Bengal (100 µl, i.v., 20 mg/ml in saline; Thermo Fisher Scientific, A17053) was injected via a tail vein catheter. The dye was then photoactivated by performing line scans along the midline of the target vessel at an excitation wavelength of 720 nm and a power of 30 mW (measured under the objective) until blood flow ceased.

### Behavioral Training for Awake Two-Photon Imaging

Training for awake imaging started once the animals had fully recovered from the craniotomy. The animals were familiarized with the experimenter and the head fixation procedure. By gradually increasing the fixation time over several days, animals became accustomed to the head fixation. Once animals were comfortable with the head fixation procedure, a water deprivation paradigm was introduced. In this paradigm, animals only received water droplets from a spout while remaining still for a specific duration. This training aimed to encourage animals to sit still during the experimental oxygen measurements, minimizing movement-induced artifacts. Most animals learned to tolerate head fixation without movement, eliminating the necessity for the water deprivation paradigm during the actual experimental imaging session.

### pO_2_ Measurements in Awake Animals

Animals trained for awake imaging were briefly anesthetized with 1.5-2% isoflurane on the baseline imaging day to inject Oxyphor 2P (10 μl, initial conc. 1.2 mM) via the cisterna magna according to a previously described protocol^37^. While under anesthesia, 50 μl of 70 kDa Texas Red dextran or QTracker655TM was additionally injected via the tail vein. Oxyphor 2P remained in the brain tissue for several days, allowing consecutive imaging sessions without needing probe reinjection^19^. After cisterna magna injection, animals were allowed to recover for a minimum of 30 min before the awake imaging started. pO_2_ measurements were performed on 4 consecutive days (pre-irradiation, post-irradiation, 24 h post, 48 h post).

A 28 × 28 point grid was positioned over the region of interest and centered on the capillary of interest. Each point in the grid was excited at 960 nm for 10 µs, and the emitted phosphorescent signal was collected for 260 μs. One excitation-collection cycle had a total duration of 270 μs, and each cycle was repeated 300 times before moving to the next point in the grid. The entire grid measuring cycle was repeated 12 times, resulting in the collection of 2600 decay points used to calculate absolute O_2_ values throughout the grid. To account for the cooling effect of two-photon imaging through a chronic cranial window using a water immersion objective, absolute pO_2_ values were calculated based on a brain temperature of 34.5 °C. A more detailed description of O_2_ data extraction and calculation can be found in previous publications^19, 37^.

### Imaging Data Analysis

All two-photon imaging data were processed using FIJI (ImageJ 2.1.0). The individual imaging channels were merged, and maximum intensity projections were generated following the specifications outlined in the figure legend. For images that were not quantitative, gamma values were adjusted non-linearly to enhance the visibility of low-intensity structures. Graphical illustrations were created using Affinity Designer (Serif Europe, UK) and Biorender (Biorender, Toronto, Canada).

### Statistical Analysis

Quantitative datasets were managed in Excel (Microsoft Corporation, Redmond, WA, USA), while statistical analyses were performed using GraphPad Prism (version 10.0; GraphPad Software, La Jolla, CA, USA). The statistical tests applied, along with the number of animals (N) and capillaries (n), are detailed in the figure legends. Data are presented as mean ± standard error of the mean (SEM). The D’Agostino-Pearson omnibus normality test was used to assess data distribution. For datasets that were found to be normally distributed, an unpaired or paired Student’s t-test was used to compare two groups. Non-normally distributed data were analyzed using the Mann-Whitney test. Comparisons involving three groups were assessed using analysis of variance (ANOVA) or, for non-parametric data, the Kruskal-Wallis test (unpaired data) or the Friedman test (paired data), with Dunn’s test applied for multiple comparisons.

### Data and Material Availability

The main text and associated source files include all relevant data supporting this study’s findings. Requests regarding the transgenic mouse lines used in this study should be directed to the principal investigator.

## ACKNOWLEDGMENTS

We thank the past and present laboratory members for their contributions to this project. We sincerely thank C. Betsholtz for the CLDN5-GFP mice. We also appreciate Lubor Borsig for providing CX3CR1-GFP mice and A. Badimon and A. Schäfer for donating clopidogrel. Thanks to S. Weber, H. Osswald, N. Binini, and A. Siebert for their support. BW was supported by the Swiss National Science Foundation (grant numbers 310030_219656 and 310030_182703). SV acknowledges the support by the U.S. National Institutes of Health through grants NS092986 and EB028941.

## AUTHOR CONTRIBUTIONS

JC, EE, CG, MTW, MA, SV and BW conceptualized and designed the study. TE, SA, TT and SV developed the photosensitizer and performed spectroscopic measurements. MV and PC performed singlet oxygen measurements. JC, EE, CG, LR, SA, TE, TT and SV contributed to data acquisition and analysis. JC, MTW, MA, SV and BW prepared the manuscript and figures with input from all authors.

## SUPPLEMENTARY FIGURES

**Supplementary Figure S1:**
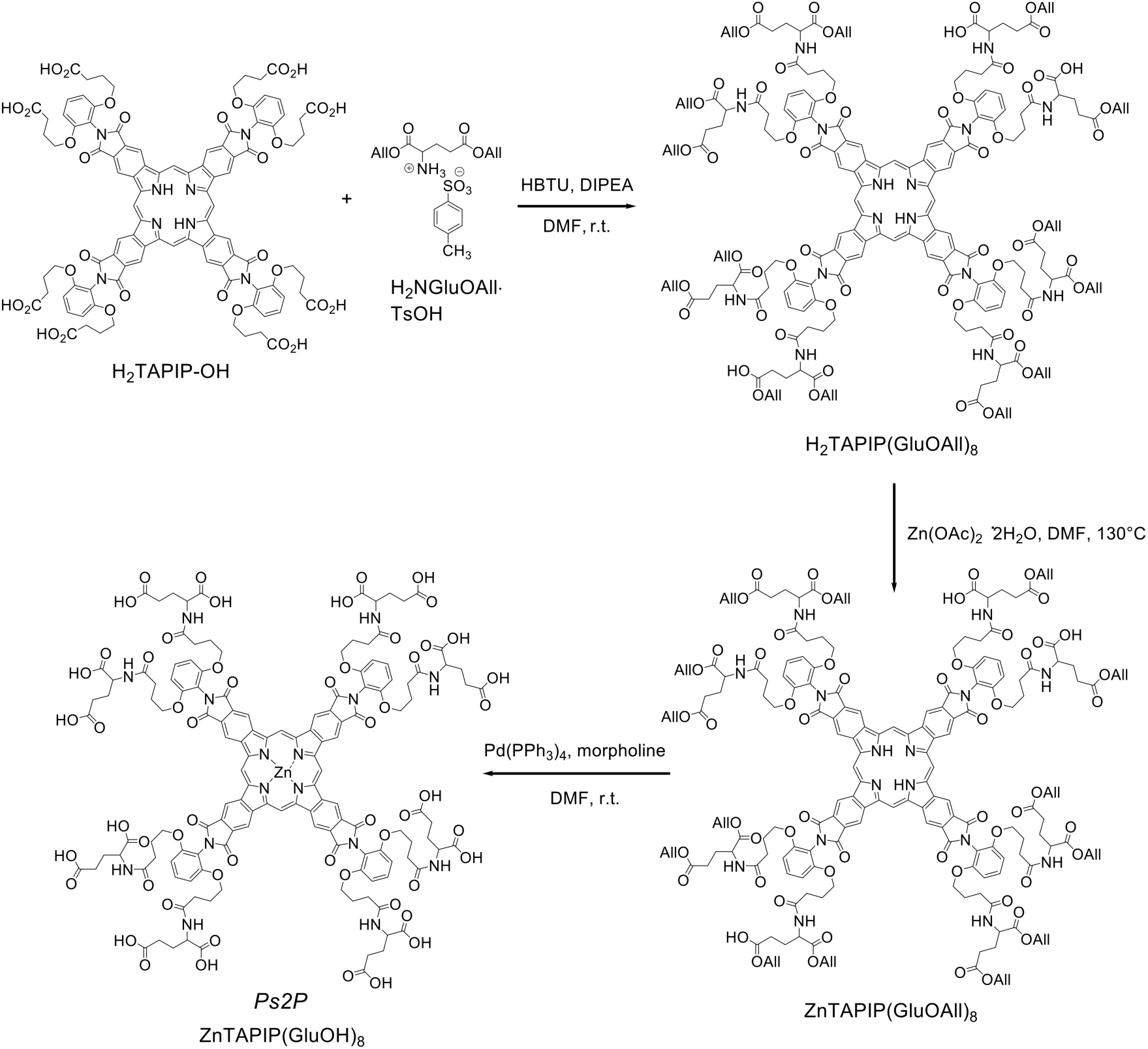
Synthesis of *PS2P*.

**Supplementary Figure S2:**
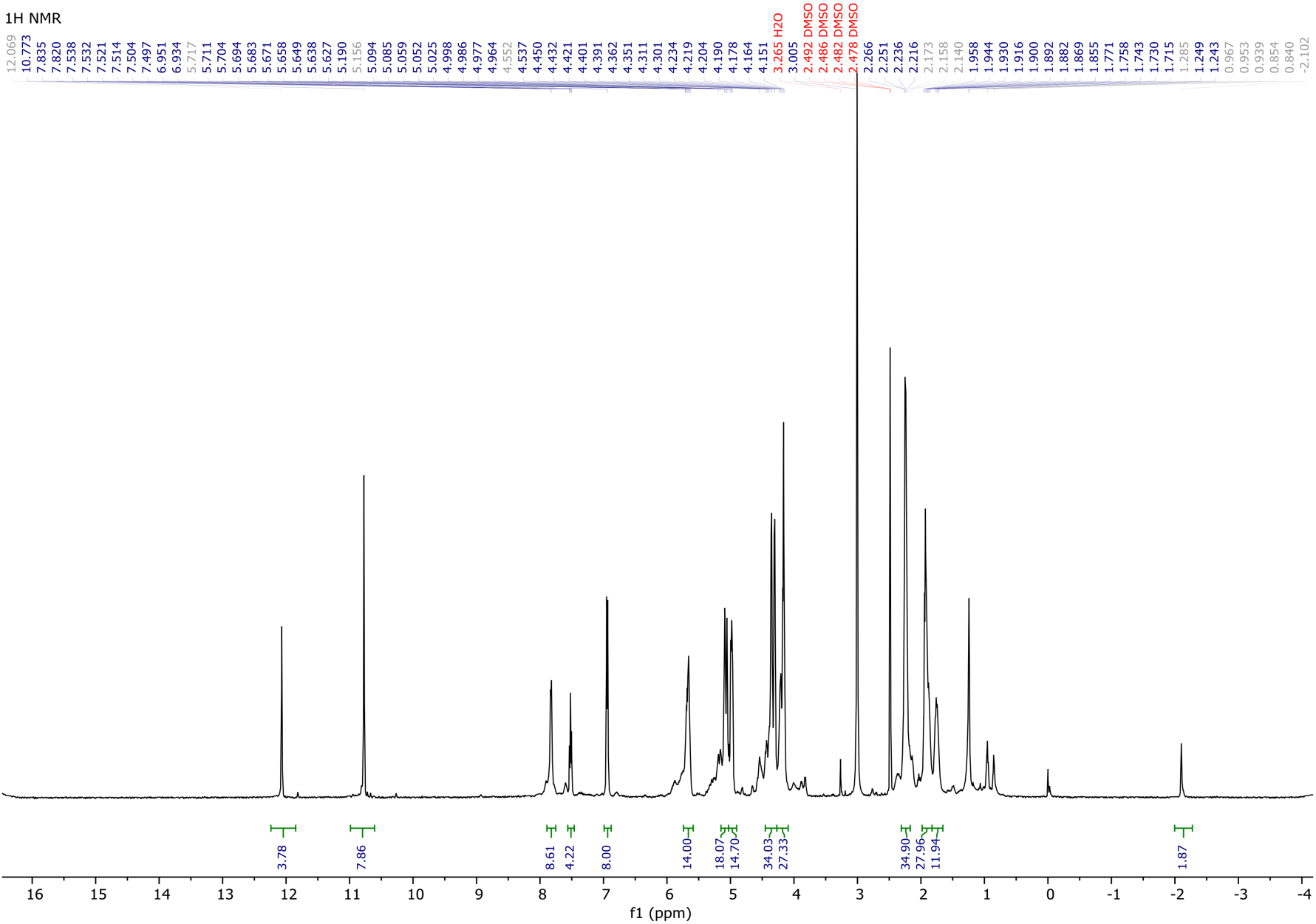
^1^H NMR spectrum of H_2_TAPIP(GluOAll)_8_.

**Supplementary Figure S3:**
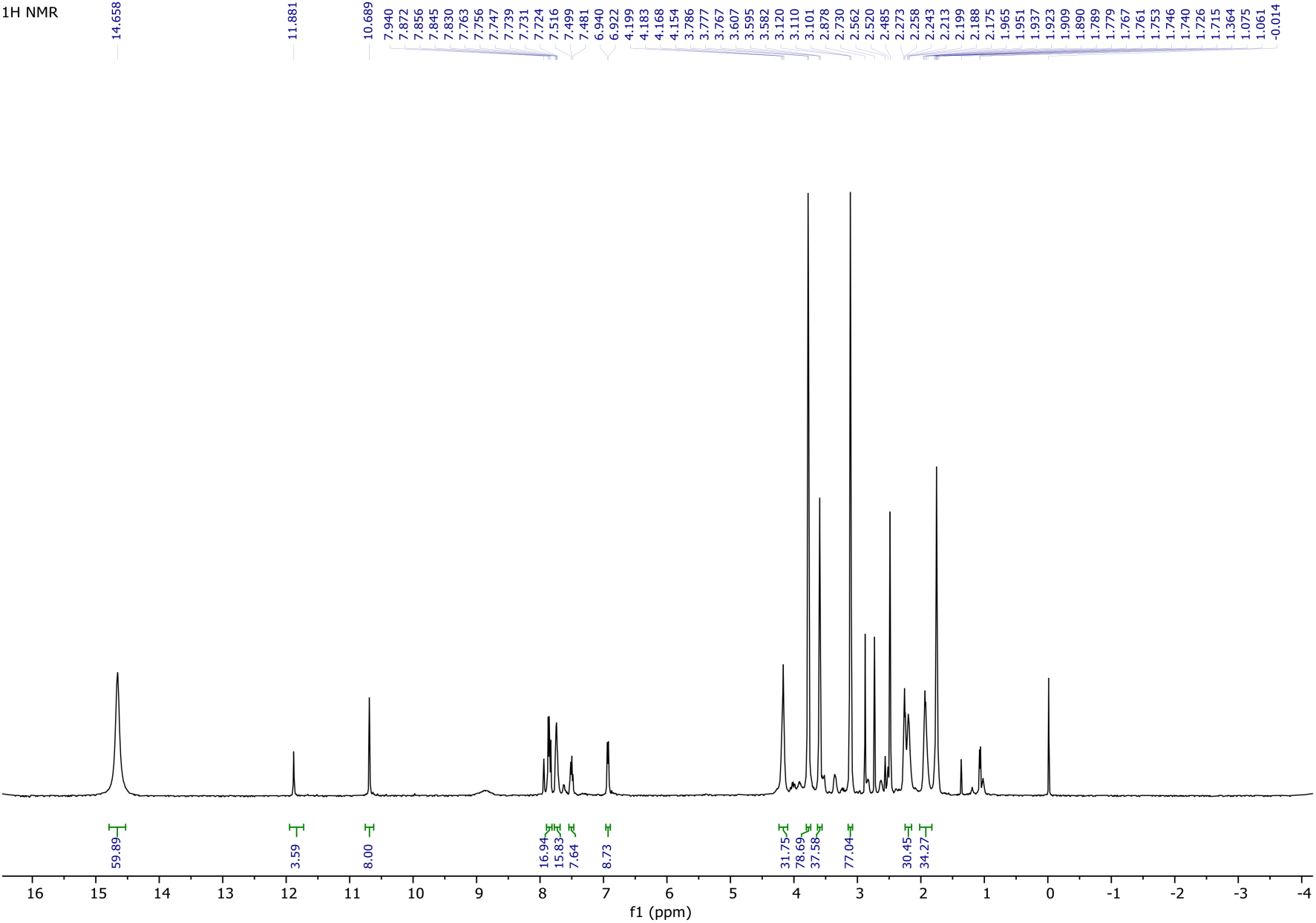
^1^H NMR spectrum of ZnTAPIP(GluOH)_8_.

## REFERENCES

1. Shih, A.Y., Blinder, P., Tsai, P.S., Friedman, B., Stanley, G., Lyden, P.D., and Kleinfeld, D. (2013). The smallest stroke: occlusion of one penetrating vessel leads to infarction and a cognitive deficit. Nat. Neurosci. 16, 55–63. 10.1038/nn.3278.

2. Nguyen, J., Nishimura, N., Fetcho, R.N., Iadecola, C., and Schaffer, C.B. (2011). Occlusion of cortical ascending venules causes blood flow decreases, reversals in flow direction, and vessel dilation in upstream capillaries. J. Cereb. Blood Flow Metab. Off. J. Int. Soc. Cereb. Blood Flow Metab. 31, 2243–2254. 10.1038/jcbfm.2011.95.

3. Nishimura, N., Schaffer, C.B., Friedman, B., Lyden, P.D., and Kleinfeld, D. (2007). Penetrating arterioles are a bottleneck in the perfusion of neocortex. Proc. Natl. Acad. Sci. U. S. A. 104, 365–370. 10.1073/pnas.0609551104.

4. Glück, C., Zhou, Q., Droux, J., Chen, Z., Glandorf, L., Wegener, S., Razansky, D., Weber, B., and El Amki, M. (2024). Pia-FLOW: Deciphering hemodynamic maps of the pial vascular connectome and its response to arterial occlusion. Proc. Natl. Acad. Sci. 121, e2402624121. 10.1073/pnas.2402624121.

5. Yoon, J.-H., Shin, P., Joo, J., Kim, G.S., Oh, W.-Y., and Jeong, Y. (2022). Increased capillary stalling is associated with endothelial glycocalyx loss in subcortical vascular dementia. J. Cereb. Blood Flow Metab. 42, 1383–1397. 10.1177/0271678X221076568.

6. Shih, A.Y., Hyacinth, H.I., Hartmann, D.A., and Van Veluw, S.J. (2018). Rodent Models of Cerebral Microinfarct and Microhemorrhage. Stroke 49, 803–810. 10.1161/STROKEAHA.117.016995.

7. van Veluw, S.J., Shih, A.Y., Smith, E.E., Chen, C., Schneider, J.A., Wardlaw, J.M., Greenberg, S.M., and Biessels, G.J. (2017). Detection, risk factors, and functional consequences of cerebral microinfarcts. Lancet Neurol. 16, 730–740. 10.1016/S1474-4422(17)30196-5.

8. El Amki, M., Glück, C., Binder, N., Middleham, W., Wyss, M.T., Weiss, T., Meister, H., Luft, A., Weller, M., Weber, B., et al. (2020). Neutrophils Obstructing Brain Capillaries Are a Major Cause of No-Reflow in Ischemic Stroke. Cell Rep. 33, 108260. 10.1016/j.celrep.2020.108260.

9. Erdener, Ş.E., Tang, J., Kılıç, K., Postnov, D., Giblin, J.T., Kura, S., Chen, I.A., Vayisoğlu, T., Sakadžić, S., Schaffer, C.B., et al. (2021). Dynamic capillary stalls in reperfused ischemic penumbra contribute to injury: A hyperacute role for neutrophils in persistent traffic jams. J. Cereb. Blood Flow Metab. 41, 236–252. 10.1177/0271678X20914179.

10. Sharma, S., Cheema, M., Reeson, P.L., Narayana, K., Boghozian, R., Cota, A.P., Brosschot, T.P., FitzPatrick, R.D., Körbelin, J., Reynolds, L.A., et al. (2024). A pathogenic role for IL-10 signalling in capillary stalling and cognitive impairment in type 1 diabetes. Nat. Metab. 6, 2082–2099. 10.1038/s42255-024-01159-9.

11. Ali, M., Falkenhain, K., Njiru, B.N., Murtaza-Ali, M., Ruiz-Uribe, N.E., Haft-Javaherian, M., Catchers, S., Nishimura, N., Schaffer, C.B., and Bracko, O. (2022). VEGF signalling causes stalls in brain capillaries and reduces cerebral blood flow in Alzheimer’s mice. Brain 145, 1449–1463. 10.1093/brain/awab387.

12. Cruz Hernández, J.C., Bracko, O., Kersbergen, C.J., Muse, V., Haft-Javaherian, M., Berg, M., Park, L., Vinarcsik, L.K., Ivasyk, I., Rivera, D.A., et al. (2019). Neutrophil adhesion in brain capillaries reduces cortical blood flow and impairs memory function in Alzheimer’s disease mouse models. Nat. Neurosci. 22, 413–420. 10.1038/s41593-018-0329-4.

13. Brown, W.R., and Thore, C.R. (2011). Review: Cerebral microvascular pathology in ageing and neurodegeneration. Neuropathol. Appl. Neurobiol. 37, 56–74. 10.1111/j.1365-2990.2010.01139.x.

14. Nelson, A.R., Sweeney, M.D., Sagare, A.P., and Zlokovic, B.V. (2016). Neurovascular dysfunction and neurodegeneration in dementia and Alzheimer’s disease. Biochim. Biophys. Acta BBA - Mol. Basis Dis. 1862, 887–900. 10.1016/j.bbadis.2015.12.016.

15. Silasi, G., She, J., Boyd, J.D., Xue, S., and Murphy, T.H. (2015). A mouse model of small-vessel disease that produces brain-wide-identified microocclusions and regionally selective neuronal injury. J. Cereb. Blood Flow Metab. Off. J. Int. Soc. Cereb. Blood Flow Metab. 35, 734–738. 10.1038/jcbfm.2015.8.

16. Wang, M., Iliff, J.J., Liao, Y., Chen, M.J., Shinseki, M.S., Venkataraman, A., Cheung, J., Wang, W., and Nedergaard, M. (2012). Cognitive deficits and delayed neuronal loss in a mouse model of multiple microinfarcts. J. Neurosci. Off. J. Soc. Neurosci. 32, 17948–17960. 10.1523/JNEUROSCI.1860-12.2012.

17. Nishimura, N., Schaffer, C.B., Friedman, B., Tsai, P.S., Lyden, P.D., and Kleinfeld, D. (2006). Targeted insult to subsurface cortical blood vessels using ultrashort laser pulses: three models of stroke. Nat. Methods 3, 99–108. 10.1038/nmeth844.

18. Taylor, Z.J., Hui, E.S., Watson, A.N., Nie, X., Deardorff, R.L., Jensen, J.H., Helpern, J.A., and Shih, A.Y. (2016). Microvascular basis for growth of small infarcts following occlusion of single penetrating arterioles in mouse cortex. J. Cereb. Blood Flow Metab. Off. J. Int. Soc. Cereb. Blood Flow Metab. 36, 1357–1373. 10.1177/0271678X15608388.

19. Esipova, T. V., Barrett, M.J.P., Erlebach, E., Masunov, A.E., Weber, B., and Vinogradov, S.A. (2019). Oxyphor 2P: A High-Performance Probe for Deep-Tissue Longitudinal Oxygen Imaging. Cell Metab. 29, 736–744.e7. 10.1016/j.cmet.2018.12.022.

20. Algorri, J.F., Ochoa, M., Roldán-Varona, P., Rodríguez-Cobo, L., and López-Higuera, J.M. (2021). Photodynamic Therapy: A Compendium of Latest Reviews. Cancers 13, 4447. 10.3390/cancers13174447.

21. Correia, J.H., Rodrigues, J.A., Pimenta, S., Dong, T., and Yang, Z. (2021). Photodynamic Therapy Review: Principles, Photosensitizers, Applications, and Future Directions. Pharmaceutics 13, 1332. 10.3390/pharmaceutics13091332.

22. Pham, T.C., Nguyen, V.-N., Choi, Y., Lee, S., and Yoon, J. (2021). Recent Strategies to Develop Innovative Photosensitizers for Enhanced Photodynamic Therapy. Chem. Rev. 121, 13454–13619. 10.1021/acs.chemrev.1c00381.

23. Bregnhøj, M., Westberg, M., Jensen, F., and Ogilby, P.R. (2016). Solvent-dependent singlet oxygen lifetimes: temperature effects implicate tunneling and charge-transfer interactions. Phys. Chem. Chem. Phys. 18, 22946–22961. 10.1039/C6CP01635A.

24. Soleimany, A., Aghmiouni, D.K., Amirikhah, M., Shokrgozar, M.A., Khoee, S., and Sarmento, B. (2024). Two-Photon Mediated Cancer Therapy: A Comprehensive Review on Two-Photon Photodynamic Therapy and Two-Photon-Activated Therapeutic Delivery Systems. Adv. Funct. Mater. 34, 2408594. 10.1002/adfm.202408594.

25. Juvekar, V., Lee, D.J., Park, T.G., Samanta, R., Kasar, P., Kim, C., Rotermund, F., and Kim, H.M. (2024). Two-photon excitation photosensitizers for photodynamic therapy: From small-molecules to nano-complex systems. Coord. Chem. Rev. 506, 215711. 10.1016/j.ccr.2024.215711.

26. Kazmi, S.M.S., Salvaggio, A.J., Estrada, A.D., Hemati, M.A., Shaydyuk, N.K., Roussakis, E., Jones, T.A., Vinogradov, S.A., and Dunn, A.K. (2013). Three-dimensional mapping of oxygen tension in cortical arterioles before and after occlusion. Biomed. Opt. Express 4, 1061–1073. 10.1364/BOE.4.001061.

27. Srivatsan, A., Missert, J.R., Upadhyay, S.K., and Pandey, R.K. (2015). Porphyrin-based photosensitizers and the corresponding multifunctional nanoplatforms for cancer-imaging and phototherapy. J. Porphyr. Phthalocyanines 19, 109–134. 10.1142/S1088424615300037.

28. Collins, H.A., Khurana, M., Moriyama, E.H., Mariampillai, A., Dahlstedt, E., Balaz, M., Kuimova, M.K., Drobizhev, M., Yang, V.X.D., Phillips, D., et al. (2008). Blood-vessel closure using photosensitizers engineered for two-photon excitation. Nat. Photonics 2, 420–424. 10.1038/nphoton.2008.100.

29. Esipova, T.V., Rivera-Jacquez, H.J., Weber, B., Masunov, A.E., and Vinogradov, S.A. (2016). Two-Photon Absorbing Phosphorescent Metalloporphyrins: Effects of π-Extension and Peripheral Substitution. J. Am. Chem. Soc. 138, 15648–15662. 10.1021/jacs.6b09157.

30. Esipova, T.V., Rivera-Jacquez, H.J., Weber, B., Masunov, A.E., and Vinogradov, S.A. (2017). Stabilizing g-States in Centrosymmetric Tetrapyrroles: Two-Photon-Absorbing Porphyrins with Bright Phosphorescence. J. Phys. Chem. A 121, 6243–6255. 10.1021/acs.jpca.7b04333.

31. Ravotto, L., Meloni, S.L., Esipova, T.V., Masunov, A.E., Anna, J.M., and Vinogradov, S.A. (2020). Three-Photon Spectroscopy of Porphyrins. J. Phys. Chem. A 124, 11038–11050. 10.1021/acs.jpca.0c08334.

32. Wilkinson, F., Helman, W.P., and Ross, A.B. (1993). Quantum Yields for the Photosensitized Formation of the Lowest Electronically Excited Singlet State of Molecular Oxygen in Solution. J. Phys. Chem. Ref. Data 22, 113–262. 10.1063/1.555934.

33. Ahn, S.J., Ruiz-Uribe, N.E., Li, B., Porter, J., Sakadzic, S., and Schaffer, C.B. (2020). Label-free assessment of hemodynamics in individual cortical brain vessels using third harmonic generation microscopy. Biomed. Opt. Express 11, 2665. 10.1364/BOE.385848.

34. Erdener, Ş.E., Tang, J., Sajjadi, A., Kılıç, K., Kura, S., Schaffer, C.B., and Boas, D.A. (2019). Spatio-temporal dynamics of cerebral capillary segments with stalling red blood cells. J. Cereb. Blood Flow Metab. Off. J. Int. Soc. Cereb. Blood Flow Metab. 39, 886–900. 10.1177/0271678X17743877.

35. Finikova, O.S., Lebedev, A.Y., Aprelev, A., Troxler, T., Gao, F., Garnacho, C., Muro, S., Hochstrasser, R.M., and Vinogradov, S.A. (2008). Oxygen Microscopy by Two-Photon-Excited Phosphorescence. ChemPhysChem 9, 1673–1679. 10.1002/cphc.200800296.

36. Sakadžić, S., Roussakis, E., Yaseen, M.A., Mandeville, E.T., Srinivasan, V.J., Arai, K., Ruvinskaya, S., Devor, A., Lo, E.H., Vinogradov, S.A., et al. (2010). Two-photon high-resolution measurement of partial pressure of oxygen in cerebral vasculature and tissue. Nat. Methods 7, 755–759. 10.1038/nmeth.1490.

37. Erlebach, E., Ravotto, L., Wyss, M.T., Condrau, J., Troxler, T., Vinogradov, S.A., and Weber, B. (2022). Measurement of cerebral oxygen pressure in living mice by two-photon phosphorescence lifetime microscopy. STAR Protoc. 3, 101370. 10.1016/j.xpro.2022.101370.

38. Császár, E., Lénárt, N., Cserép, C., Környei, Z., Fekete, R., Pósfai, B., Balázsfi, D., Hangya, B., Schwarcz, A.D., Szabadits, E., et al. (2022). Microglia modulate blood flow, neurovascular coupling, and hypoperfusion via purinergic actions. J. Exp. Med. 219, e20211071. 10.1084/jem.20211071.

39. Davalos, D., Grutzendler, J., Yang, G., Kim, J.V., Zuo, Y., Jung, S., Littman, D.R., Dustin, M.L., and Gan, W.-B. (2005). ATP mediates rapid microglial response to local brain injury in vivo. Nat. Neurosci. 8, 752–758. 10.1038/nn1472.

40. Nimmerjahn, A., Kirchhoff, F., and Helmchen, F. (2005). Resting Microglial Cells Are Highly Dynamic Surveillants of Brain Parenchyma in Vivo. Science 308, 1314–1318. 10.1126/science.1110647.

41. Condrau, J., Glück, C., Wyss, M.T., Hösli, L., Erlebach, E., Zanker, H., Ravotto, L., von Faber-Castell, A., Esipova, T.V., Herwerth, M., et al. (2025). Intrinsic Endothelial Remodeling Enables Cerebral Microvascular Repair. bioRxiv.

42. Delafontaine-Martel, P., Zhang, C., Lu, X., Damseh, R., Lesage, F., and Marchand, P.J. (2023). Targeted capillary photothrombosis via multiphoton excitation of Rose Bengal. J. Cereb. Blood Flow Metab. 43, 1713–1725. 10.1177/0271678X231151560.

43. Fukuda, M., Matsumara, T., Suda, T., and Hirase, H. (2022). Depth-targeted intracortical microstroke by two-photon photothrombosis in rodent brain. Neurophotonics 9. 10.1117/1.NPh.9.2.021910.

44. Xi, W., Zhu, L., Zhang, H., Wang, M., and Roe, A. (2024). Single microvessel occlusion technology (PLP) produces cortical microvasculature redistribution and neuronal network functional adaptation. In Optogenetics and Optical Manipulation 2024 (SPIE), p. 128290J. 10.1117/12.3012512.

45. Sinks, L.E., Robbins, G.P., Roussakis, E., Troxler, T., Hammer, D.A., and Vinogradov, S.A. (2010). Two-Photon Microscopy of Oxygen: Polymersomes as Probe Carrier Vehicles. J. Phys. Chem. B 114, 14373–14382. 10.1021/jp100353v.

46. Schmid, F., Conti, G., Jenny, P., and Weber, B. (2021). The severity of microstrokes depends on local vascular topology and baseline perfusion. eLife 10, e60208. 10.7554/eLife.60208.

47. Farkas, E., and Luiten, P.G.M. (2001). Cerebral microvascular pathology in aging and Alzheimer’s disease. Prog. Neurobiol. 64, 575–611. 10.1016/S0301-0082(00)00068-X.

48. Madsen, L.S., Parbo, P., Ismail, R., Gottrup, H., Østergaard, L., Brooks, D.J., and Eskildsen, S.F. (2023). Capillary dysfunction correlates with cortical amyloid load in early Alzheimer’s disease. Neurobiol. Aging 123, 1–9. 10.1016/j.neurobiolaging.2022.12.006.

49. Nortley, R., Korte, N., Izquierdo, P., Hirunpattarasilp, C., Mishra, A., Jaunmuktane, Z., Kyrargyri, V., Pfeiffer, T., Khennouf, L., Madry, C., et al. (2019). Amyloid β oligomers constrict human capillaries in Alzheimer’s disease via signaling to pericytes. Science 365, eaav9518. 10.1126/science.aav9518.

50. Iadecola, C. (2017). The Neurovascular Unit Coming of Age: A Journey through Neurovascular Coupling in Health and Disease. Neuron 96, 17–42. 10.1016/j.neuron.2017.07.030.

51. Plunkett, S., Khatib, M.E., Sencan, I., E. Porter, J. N. Kumar, A.T., E. Collins, J., Sakadžić, S., and A. Vinogradov, S. (2020). In vivo deep-tissue microscopy with UCNP/Janus-dendrimers as imaging probes: resolution at depth and feasibility of ratiometric sensing. Nanoscale 12, 2657–2672. 10.1039/C9NR07778B.

52. Würth, C., Grabolle, M., Pauli, J., Spieles, M., and Resch-Genger, U. (2013). Relative and absolute determination of fluorescence quantum yields of transparent samples. Nat. Protoc. 8, 1535–1550. 10.1038/nprot.2013.087.

53. Makarov, N.S., Drobizhev, M., and Rebane, A. (2008). Two-photon absorption standards in the 550-1600 nm excitation wavelength range. Opt. Express 16, 4029–4047. 10.1364/OE.16.004029.

54. Mayrhofer, J.M., Haiss, F., Haenni, D., Weber, S., Zuend, M., Barrett, M.J.P., Ferrari, K.D., Maechler, P., Saab, A.S., Stobart, J.L., et al. (2015). Design and performance of an ultra-flexible two-photon microscope for in vivo research. Biomed. Opt. Express 6, 4228. 10.1364/BOE.6.004228.

55. Pologruto, T.A., Sabatini, B.L., and Svoboda, K. (2003). ScanImage: Flexible software for operating laser scanning microscopes. Biomed. Eng. OnLine 2, 13. 10.1186/1475-925X-2-13.

56. Barrett, M.J.P., Ferrari, K.D., Stobart, J.L., Holub, M., and Weber, B. (2018). CHIPS: an Extensible Toolbox for Cellular and Hemodynamic Two-Photon Image Analysis. Neuroinformatics 16, 145–147. 10.1007/s12021-017-9344-y.

57. Drew, P.J., Blinder, P., Cauwenberghs, G., Shih, A.Y., and Kleinfeld, D. (2010). Rapid determination of particle velocity from space-time images using the Radon transform. J. Comput. Neurosci. 29, 5–11. 10.1007/s10827-009-0159-1.

58. Binder, N.F., El Amki, M., Glück, C., Middleham, W., Reuss, A.M., Bertolo, A., Thurner, P., Deffieux, T., Lambride, C., Epp, R., et al. (2024). Leptomeningeal collaterals regulate reperfusion in ischemic stroke and rescue the brain from futile recanalization. Neuron 112, 1456–1472.e6. 10.1016/j.neuron.2024.01.031.

